# Single-cell phenotyping and RNA sequencing reveal novel patterns of gene expression heterogeneity and regulation during growth and stress adaptation in a unicellular eukaryote

**DOI:** 10.1101/306795

**Authors:** Malika Saint, François Bertaux, Wenhao Tang, Xi-Ming Sun, Laurence Game, Anna Köferle, Jürg Bähler, Vahid Shahrezaei, Samuel Marguerat

**Affiliations:** MRC London Institute of Medical Sciences (LMS), Du Cane Road, London W12 0NN, UK; Institute of Clinical Sciences (ICS), Faculty of Medicine, Imperial College London, Du Cane Road, London W12 0NN, UK; Department of Mathematics, Faculty of Natural Sciences, Imperial College, London SW7 2AZ, UK; University College London, Research Department of Genetics, Evolution & Environment and UCL Cancer Institute, London, WC1E 6BT, UK

## Abstract

Cell-to-cell variability is central for microbial populations and contributes to cell function, stress adaptation and drug resistance. Gene-expression heterogeneity underpins this variability, but has been challenging to study genome-wide. Here, we report an integrated approach for imaging of individual fission yeast cells followed by single-cell RNA sequencing (scRNA-seq) and novel Bayesian normalisation. We analyse >2000 single cells and >700 matching RNA controls in various environmental conditions and identify sets of highly variable genes. Combining scRNA-seq with cell-size measurements provides unique insights into genes regulated during cell growth and division in single cells, including genes whose expression does not scale with cell size. We further analyse the heterogeneity and dynamics of gene expression during adaptive and acute responses to changing environments. Entry into stationary phase is preceded by a gradual, synchronised adaptation in gene regulation, followed by highly variable gene expression when growth decreases. Conversely, a sudden and acute heat-shock leads to a stronger and coordinated response and adaptation across cells. This analysis reveals that the extent and dynamics of global gene-expression heterogeneity is regulated in response to different physiological conditions within populations of a unicellular eukaryote. In summary, this works illustrates the potential of combined transcriptomics and imaging analysis in single cells to provide comprehensive and unbiased mechanistic understanding of cell-to-cell variability in microbial communities.

---

Expression of DNA into mRNA and proteins is a fundamental biological process that governs most aspects of the cell physiology. Gene expression is tightly regulated at multiple levels, including chromatin structure, transcription, and post-transcriptional processes such as mRNA stability and translation. This multi-layered process underpins robust and timely expression of single proteins as well as coordinated regulation of entire genetic programmes that include dozens of genes. Yet, even in relatively constant environments, expression of specific genes vary between genetically identical cells, giving rise to cell-to-cell heterogeneity in mRNA numbers and concentrations^1–3^. Cell-to-cell variability in gene expression results from different phenomena. The random timing of biological reactions makes the transcription process stochastic. This form of variability, also called intrinsic noise, is gene specific and depends on promoter sequence and chromatin states^45^. Heterogeneity in quantitative cellular traits such as size, growth rate, or concentration of general transcription factors also shapes gene expression variability across cells in complex, non-trivial ways. This form of variability is not entirely stochastic and depends on other single cell attributes that affect biomolecule numbers^6,7^. Cells adopt distinct states characterised by transcriptional programmes regulating the concentration of specific mRNAs. Examples are the progression through the different phases of the cell cycle or of adopting different, cycling metabolic states^8^. Different states can co-exist in cell populations or in complex tissues leading to dynamic, yet deterministic, cell-to-cell variability in gene expression. Finally, cells in complex tissues feature diverse differentiated states that are important for tissue architecture and function. Although reversible and plastic, this form of individuality is not expected to be as dynamic as more transient cellular states.

The development of RNA sequencing protocols that support the analysis of entire transcriptomes from single cells has been instrumental for our understanding of cell-to-cell variability and phenotypic heterogeneity in multicellular organisms^9^. Because such approaches sample expression levels of large numbers of genes in an unbiased manner, they can provide insights into the molecular complexity of healthy tissues and tumours, and reveal novel and rare types of cells, affording better understanding of tissue biology in health and disease^10–12^ Critically, the development of comparable approaches for sampling gene-expression heterogeneity in microbial cells has lagged behind, mostly due to the small size and resistant cell walls of these organisms^13^. Yet, phenotypic and gene-expression variability between single unicellular organisms in a population is conceptually very different from the heterogeneity observed in metazoan tissues. Moreover, determining the extent of cell-to-cell variability in gene regulation in microbial populations is of central importance to our understanding of resistance to antibiotics, cellular adaptation or population dynamics and evolution^13,14^.

Here, we overcome these limitations and develop an integrated experimental and computational framework to image individual cells of fission yeast (*Schizosaccharomyces pombe*), followed by singlecell RNA-seq analysis (scRNA-seq). Using this approach, we obtained a unique account of the heterogeneity in gene expression and cellular states as a function of cell size, growth and adaptation in a popular unicellular model organism.

## Imaging and transcriptome analysis of single fission yeast cells

We have developed an integrated pipeline for imaging and isolation of single cells into PCR tubes, using a tetrad dissection microscope, followed by transcriptome analysis by scRNA-seq (Fig. 1a, **Supplementary Notes**). With this innovative approach, we generated combined datasets consisting of microscope images and RNA-seq libraries for 2028 single cells from a range of environmental conditions, together with 780 matching control samples consisting of 3pg of total RNA (ctrRNA, **Supplementary Table S1 and S2**).

**Figure 1:**
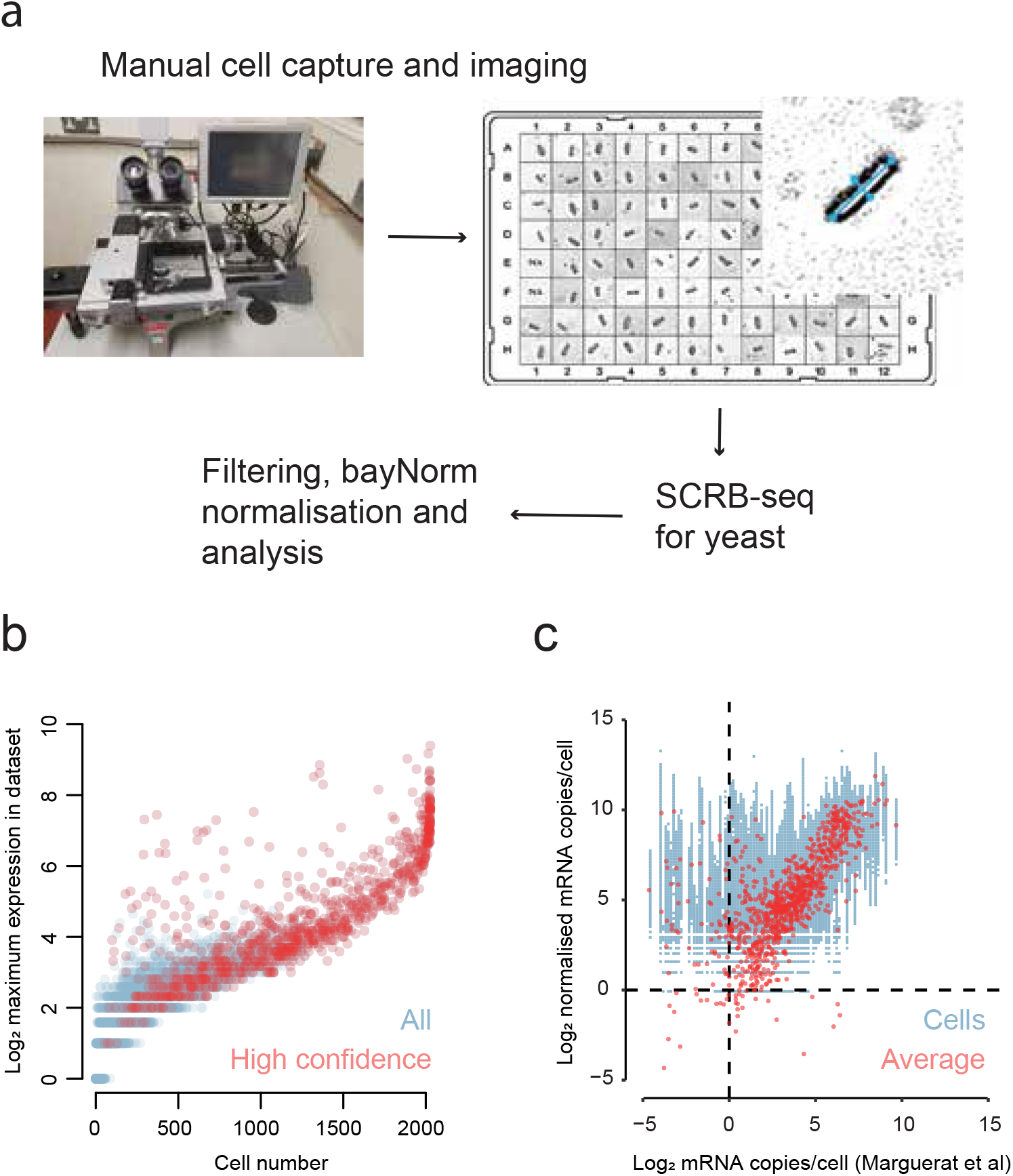
Imaging and transcriptome analysis of single fission yeast cells. **(a)** Experimental and analysis pipelines. Batches of 96 single cells are isolated and imaged using a dissection microscope. Single cell RNA-seq libraries are generated using SCRB-seq for yeast followed by filtering and normalisation using bayNorm, a novel Bayesian approach for true count inference of scRNA-seq data^20^. Processed data are used for functional and differential expression analyses. **(b)** Overall coverage of scRNA-seq datasets and selection of high-confidence genes. Maximum raw counts for each coding and non-coding mRNA across 21 datasets (2028 cells) are plotted as a function of the number of cells in which the mRNA was detected. All genes (blue) and the high-confidence genes used for all further analysis in this study (red) are shown. **(c)** Gene expression levels in single cells. Normalised scRNA-seq counts are plotted as a function of population average mRNA copies/cell data from Marguerat et al^21^. Normalised counts in single cells (blue), and average counts across cells (red) are shown. R_pearson_ = 0.6 for average counts.

RNA sequencing analysis of single yeast cells is particularly challenging because their cell wall is resistant to the commonly used lysis conditions that preserve RNA integrity. Moreover yeast transcriptomes are extremely plastic and respond to changes in external conditions within minutes making cell isolation and manipulation a source of artefacts. We have overcome both hurdles by i) snap freezing cells immediately after harvesting of the culture, a procedure that fixes and preserves both cell morphology and transcriptomes; ii) establishing a protocol for yeast cells lysis at high temperature in conditions that protects RNA integrity, bypassing the need for enzymatic digestion of the cell wall (Supplementary Notes).

To prepare sequencing libraries from the minute amounts of mRNA present in yeast cells, we have developed a variation of the SCRB-seq protocol^15,16^. A recent comparative analysis of scRNA-seq protocols has identified SCRB-seq as an accurate an efficient method^17^. Our approach targets 3’-end cDNA sequences and includes unique molecular identifiers (UMI), and benefits, in addition, from optimisations of the Smart-seq2 approach (**Supplementary Notes**)^15,16,18^. Using this method, we generated 8.2 × 10^5^ mappable sequencing reads per single cell on average, corresponding to 6721.3 unique mRNA molecules on average (Fig. S1a-c, **Supplementary Notes**, **Supplementary Table S1**). This represents a mean transcriptome coverage 〈β〉 of ~1.5% or ~8%, depending on whether spike-in controls or single-molecule fluorescence in situ hybridisation (smFISH) measurements of mRNA levels were used for calibration, respectively (Fig. S1d, **Supplementary Notes**).

With this level of coverage, we could detect 1421.1 genes per cell on average (**Supplementary Table S2**). Genes with low scRNA-seq molecule counts carried little information on real expression levels. We therefore focused this study on a consolidated list of 1011 high-confidence genes with robust measurements, which included 18.5% and 4.3% of the fission yeast coding and non-coding transcriptomes respectively (Fig. 1b, **Supplementary Notes**, **Supplemetary Table S1 and S3**). As expected, these genes were biased towards high expression but were involved in most cellular processes, showing constitutive as well as condition-specific regulation (Fig. 1b, Fig. S1e, **Supplemetary Table S5**).

Shallow transcriptome coverage is an inherent property of scRNA-seq datasets that affects their structure and makes data normalisation challenging^19^. To account for this issue, in particular for the relative impact of technical sampling noise that increases at low transcripts counts, we developed a novel Bayesian normalisation approach called bayNorm that computes estimates of true mRNA counts from raw scRNA-seq data^20^. This method relies on estimates of each cell transcriptome coverage β, also called capture efficiency, and on gene-specific priors based on pooling of the data across cells using an empirical Bayes methodology (**Supplementary Notes**)^20^. The bayNorm approach performs both normalisation and imputation of missing values, reducing drop-out levels and experimental batch effects, correcting efficiently for cell-to-cell variability in coverage, and recovering expression distributions similar to population RNA-seq and smFISH data (Fig. S1f-g, Fig. S2e, **Supplementary Table S4**)^20^. Normalised expression counts showed a good correlation with expression levels derived from cell populations^21^, while preserving information about cell-to-cell variability in mRNA expression levels (average R_pearson_ =0.78, Fig. 1c, Fig. S1h).

In summary, we developed a versatile and powerful approach for genome-wide analysis of transcriptomes in single yeast cells, while enabling integration with phenotypic features. Using this method, we generated the first combined dataset of transcriptomes and microscopy images from single yeast cells that is analysed in detail below.

## Cell-to-cell variability of transcriptome regulation

We first used our scRNA-seq data to define the repertoire of expressed genes with strong cell-to-cell variability in mRNA levels (Highly Variable Genes, HVG). Large levels of variability in mRNA expression were immediately apparent from single-cell transcriptomes (Fig. 1c). However, because of the minute amount of starting material scRNA-seq relies on, part of this variability results from random technical inaccuracies during sample preparation rather than from biology. As previously reported and expected from stochastic gene expression models^22,23^, we observed a negative relation between means and coefficient of variation (σ^2^/m, CV) of bayNorm normalised counts (Fig. S2a). Taking this relation into account, we looked for mRNAs with CVs significantly higher than the average trend in each individual cell and ctrRNA dataset collected during rapid cell proliferation when compared to simulated synthetic controls containing no biological variability (864 cells, **Supplementary Table S1**, **Supplementary Notes**). We identified 411 HVG as significant in at least one dataset (Fig. 2a, **Supplementary Notes**), of which 112 genes were also present in at least one ctrRNA dataset and were discarded as false positives (FP). This analysis resulted in 299 high-confidence HVGs (Fig. S2b, **Supplementary Table S5**). Transcript number distributions of a subset of HVGs analysed by single-molecule fluorescence *in situ* hybridisation (smFISH) confirmed their high variability (Fig. 2b, Fig. S2c-d, and **Supplementary Notes**).

**Figure 2:**
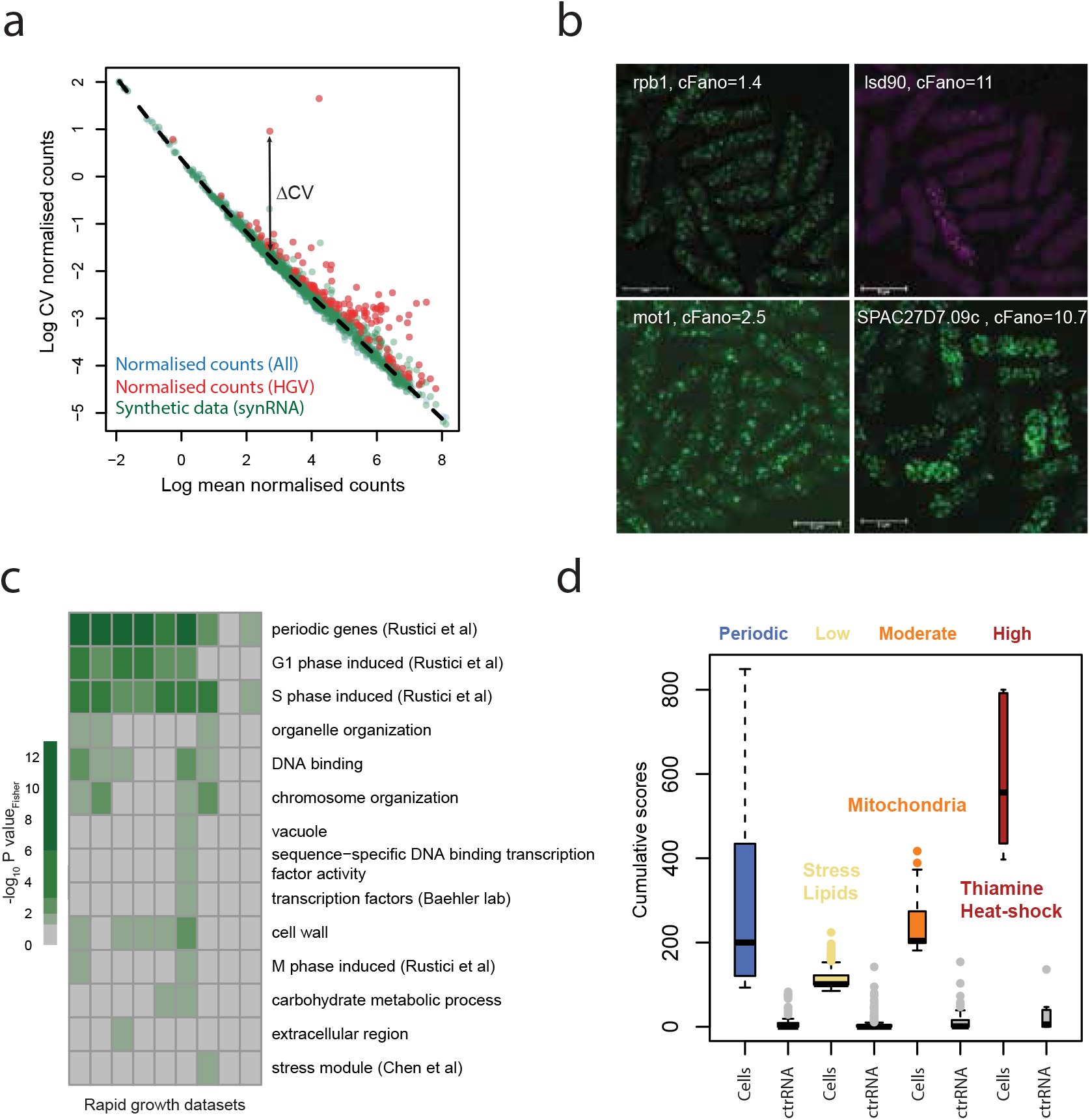
Cell-to-cell variability of fission yeast transcriptome. **(a)** Identification of highly variable genes (HVG). Coefficients of variation of normalised counts are plotted for all mRNAs (blue, mostly hidden), mRNAs called variable (red), and simulated synthetic controls used for calling variability (green, synRNA; **Supplementary Notes**) are plotted as a function of their respective mean normalised expression. Loess fit for synthetic controls is shown as a dotted line. An mRNA is called highly variable if it deviates significantly from the distribution of synthetic controls of similar mean expression. The ΔCV of an example HVG is highlighted with an arrow. **(b)** Validation by smFISH of mRNA called variable from scRNA-seq data. Representative smFISH images are shown for low-variability control *rbp1* mRNA and three mRNAs with different levels of variability (*lsd90, mot1*, and SPAC27D7.09c, see also Fig. S2c-d). Size corrected Fano factors are indicated in each plot. **(c)** Functional analysis of variable genes from 9 datasets of 96 cells during rapid proliferation. Significance of overlap of variable mRNAs in each dataset with selected functional categories is shown (-log_10_ of P_Fisher_ for overlap of HVGs from each dataset and each category). False positive mRNAs called from total RNA control experiments have been filtered out (Fig. S2b). Note that some categories are more pervasively variable across datasets than others. **(d)** Functional analysis of non-periodic HVGs. HVGs that have not been observed as cell-cycle regulated in cell populations were sorted into three categories with “low”, “moderate”, or “high” pervasive variable expression. Selected distinctive gene functions are shown in the plot.

Several studies have identified genes that are periodically expressed during the cell cycle in synchronised cell populations^24–27^. We hypothesised that those genes should be overrepresented among HVGs, as we sampled asynchronous cultures, which combine cells from different cell cycle stages. As expected, 53.3% of the top-500 periodic genes^27^ found in our datasets were HVGs (P_Fisher_ = 5.1e^-9^). Moreover, genes from functional categories associated with phase-specific expression during the cell cycle were enriched among HVGs in most datasets (Fig. 2c). These findings, and the fact that only few periodic genes were called false positives, confirms the specificity of our experimental and analysis protocols (Fig. S2b, **Supplementary Table S5**). This analysis validates our approach further and demonstrates that periodic gene expression as a function of the cell cycle is a single-cell feature of asynchronous populations and not a technical artefact of cell-cycle synchronisation^28^.

Nevertheless, most HVGs were not cell-cycle periodic genes and could not have been identified in cell populations. These HVGs were enriched for the Core Environmental Stress Response programme (CESR)^29^ and for genes related to vacuole biology, and were detected in fewer datasets than the periodic genes (Fig. 2c, Fig. S2b). This result suggests subtle gene regulation to fluctuations in external conditions in line with a recent scRNA-seq report in budding yeast^13^. Interestingly, orthologues of the 235 HVGs that had not been described as periodic were also significantly enriched in the top-100 most variable budding yeast genes (P_Fisher_ = 0. 02). This finding indicates that the architecture of gene-expression variability is at least partially conserved between the two yeasts^22^. Moreover, these HVGs were significantly over-represented among fission yeast genes most variable in response to environmental and genetic perturbations in cell populations (P_Fisher_ = 0.03)^22,30^. This finding suggests that noisy transcription of stress-responsive genes could reflect a bet-hedging mechanism to facilitate population survival to rapid, unpredictable challenges^31^. Variably expressed genes have been reported to also evolve rapidly^32,33^. Accordingly, these genes also showed significantly higher evolution rates between orthologues of 4 fission yeast species (P_Wilcox_ = 0.03)^34^.

We then split the non-periodic HVGs in three categories with increasingly pervasive variability across datasets (Fig. 2d, **Supplementary Table S5**). Moderately and highly pervasive HVGs were related to the biology of mitochondria and heat-shock proteins. The genes encoding Nmt1 and the associated biosynthetic enzyme Thi2 were also among the most pervasively variable HVGs, suggesting heterogeneity in vitamin B1 metabolism. In terms of gene-expression regulators, the transcription factor

Fil1, controlling the amino-acid starvation response, as well as the TATA-binding associated factor Mot1, a general transcription factor, were also pervasively variable (Fig 2b, S2c-d)^35,36^. The later observation is consistent with TATA box sequences being associated with more variable genes^5,32,33^. Lowly pervasive HVGs span diverse functions related to membrane biology and adaptation to external conditions. Notably, some lowly pervasive genes, like *Isd90*, could show very strong amplitudes of regulation (Fig. 2b, S2c-d). In summary, we show that genome-expression variability of proliferating fission yeast cells measured by scRNA-seq features different levels of pervasiveness, with only a subset of the variable genes reflecting periodic regulation during the cell cycle.

## Cell size dependence of transcriptome regulation

If coupled with other single-cell analytical techniques, based on imaging for instance, scRNA-seq can probe changes in gene expression globally and as a function of quantitative cell phenotypes. However, developing such combined approaches is challenging and only few examples are available to date^6,37^. Our experimental pipeline uniquely harnesses this potential as it includes microscopy images of each cell matched to their respective transcriptomes (Fig. 1a).

Using cell-length measurements extracted from images across all growth datasets, we first examined changes in global properties of scRNA-seq expression measurements as a function of cell size (**Supplementary Table S2**). The mean cell length was of 10.9μm, consistent with previously reported data for fission yeast growing in EMM2 medium. Average normalised RNA-seq counts were constant across the size range, consistent with bayNorm returning size-corrected absolute molecule numbers (which is proportional to concentrations; see **Supplementary Notes**, Fig. 1a).

When grown in EMM, fission yeast elongates exclusively during G2 phase of the cell cycle, which accounts for over two thirds of the cell-cycle time. Upon entry into mitosis, cells arrest and stop growing until after cell division, which occurs coincident with G1 and S phases. As a proof of principle, we therefore asked whether we could assign transcriptome signatures of the M/G1/S phases to large cells. As expected, large cells featured increased transcriptome fractions related to processes specific to G1/S transition and cell-wall biogenesis (Fig. 3b, **Supplementary Notes**). This observation was confirmed by visual inspection of the data (Fig. S3a). Besides cell-cycle signatures, some large cells featured increased transcriptome fractions functioning in respiratory metabolism, which is thought to occur during the reductive building phase of the yeast metabolic cycle (Fig. 3b, see below)^38^.

**Figure 3:**
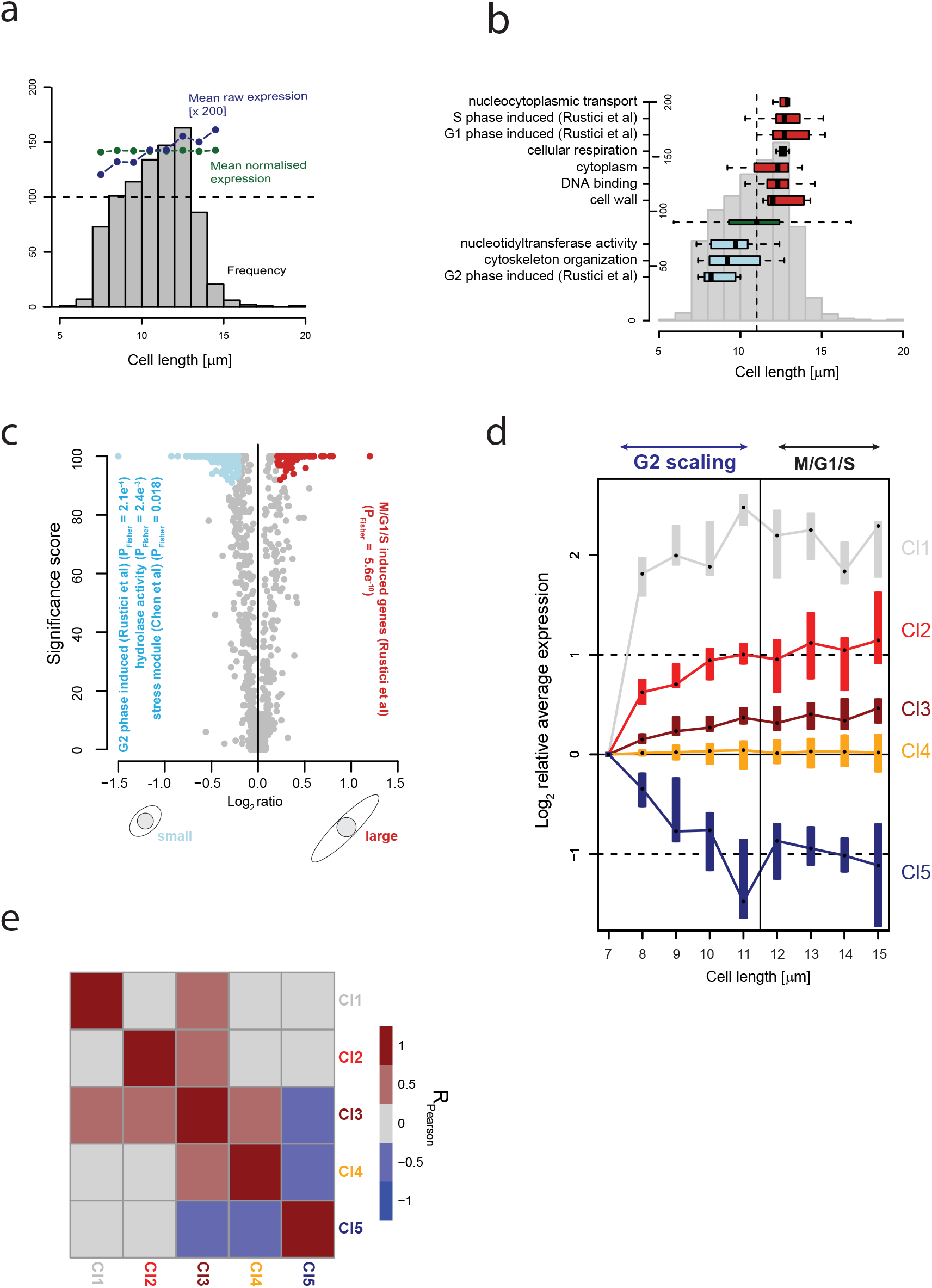
Cell-size dependence of fission yeast transcriptome. **(a)** Cell length distribution across 864 cells during rapid proliferation and global characteristics of the corresponding transcriptomes. Average raw expression scores (blue) or average bayNorm normalised expression scores (green) are shown for cell length bins of 1μm. Note the positive correlation of raw scores with cell size that is lost after normalisation. **(b)** Single cells were assigned to functional categories based on their relative transcriptome signatures. Shown are cells in categories associated with cells significantly smaller (blue) or larger (red) than the overall population size distribution (green). Data are overlaid onto the cell size frequency histogram shown in **(a)**. **(c)** Differential expression (DE) analysis between large (13-16μm) and small (8-10μm) cells. Number of bootstrap iteration showing significant DE call are plotted for each gene (P_MAST_ <0.05, total iterations = 100) as a function of MAST log_2_ DE ratios. Genes significantly induced in small and large cells are highlighted in blue and red, respectively (Supplementary table S2). Selected functional categories that are significantly enriched are shown along with enrichment P_Fisher_. **(d)** The mRNAs that change in concentration during the G2 growth phase (non-scaling genes, NSG). Average expression scores among cells shorter than 11μm in length were computed for bins of 1μm, normalised to the smallest size bin and used for k-means clustering. Genes with significant linear correlation of bayNorm normalised counts with cell size were used (P <0.05). **(e)** Co-regulation of NSG clusters. Pearson correlations between clusters from **(d)** are shown. Note that Clusters 1 and 2 are co-regulated while Clusters 3 and 5 are anti-regulated.

We then used cell-length measurements to guide our analysis of the transcriptional programmes associated with cell growth and the cell cycle. To do so, we looked for genes differentially expressed between large cells in M/G1/S phases and small, recently born cells in G2 phase using bayNorm priors specific for the two groups (Fig. 3c, Fig. S3b-c, **Supplementary Notes**). We identified 92 genes significantly up-regulated in large cells (**Supplementary Table S5**). Consistent with larger cells being in the M/G1/S phases of the cell cycle, 28.3% of these 92 genes were also periodically expressed in synchronised cell populations^24–27^. Around twice as many genes (193) were induced in small, recently born cells, of which 19.2% had been described as periodically expressed in synchronised cell populations^27^. A significant proportion of these genes belonged to the stress response programme (P_Fisher_ = 0.02) and/or had hydrolase activity (P_Fisher_ = 0.002). Large cells, on the other hand, induced several genes functioning in membrane transport involved in mitochondrial biology. These observations, together with the analysis from Fig. 3b, raised the possibility of a fission yeast metabolic cycle (YMC) in single asynchronous cells^39^. We therefore analysed gene signatures of the three proposed YMC phases: OX (oxidative), RB (reductive building) and RC (reductive charging). We found signatures of compartmentalisation with cell size and the cell cycle (Fig. S3d)^38^. RC genes were highly expressed in smaller cells, while RB genes were higher expressed in large cells when DNA replication occurs^8,38^. Altogether, this analysis demonstrates the existence of a fission yeast metabolic cycle which is synchronised with the cell cycle in single cells during rapid proliferation^40^.

The numbers of most mRNAs increase co-ordinately, or scale, with cell size in order to maintain transcript concentrations^41–43^. Accordingly, average UMI corrected raw mRNA molecule numbers per cell showed a weak but significant correlation with cell size in our data (Fig. 3a). Genes that escape this general trend and change expression as a function of cell size have not been characterised globally, yet they could represent important players in the regulation of growth and cell-size homeostasis^44^. To identify genes that escape scaling, we analysed G2 cells between birth and 11μm in size, beyond which we found strong signatures of the M/G1/S programmes (Fig. S3a, **Supplementary Notes**, **Supplementary Table S5**). We observed 78 genes that significantly and coordinately changed in concentration with cell size during G2 (P_pearson_ < 0.05). Using unsupervised k-means clustering, we defined 5 small gene clusters, 3 of which increased (Cl1-3) and 1 decreased (Cl5) in concentration (Fig. 3d, **Supplementary Notes**, **Supplementary Table S5**). We then assessed whether these clusters defined one or more cellular state by looking at their correlations between single cells (**Supplementary Notes**). We found evidence for the existence of two states in our datasets. The first was defined by Cl1 and Cl2 that were positively correlated and contained genes that are also upregulated during meiotic differentiation (Fig. 3e)^45^. Cl3 and Cl5 were anti-correlated in single cells and defined a second state containing a small number of genes functioning in carbohydrate metabolism (Fig. 3e). Although not significant, this enrichment could hint at a gradual change in cellular energy metabolism coordinated with cell-size increase. We used an orthogonal Random Forest approach to validate this analysis and confirmed that 69 of the 100 genes with the strongest non-linear correlation with cell size were also either differentially expressed between big and small cells, or escaped scaling (**Supplementary Notes**, **Supplementary table S2**). Overall, this analysis uncovered a novel form of gene expression regulation, which occurs during cell growth in G2 phase and is co-ordinated with cell size.

In summary, we find that 45.5% of the HVG genes detected in this study are either overexpressed in large M/G1/S cells or in small G2 cells, or are regulated during cell growth. Their variability is therefore not stochastic *per se* but can be understood in the light of two physiological variables, the size of the cell and its cell-cycle stage. Moreover, our data suggest the existence of a novel layer of heterogeneity occurring during the growth phase in G2. Going beyond fission yeast biology, this analysis demonstrates the unique potential of analysing scRNA-seq datasets in the context of quantitative cellular features such as cell size.

## Transcriptome heterogeneity within cell populations in response to environmental changes

Defining the impact of external signals and environmental conditions on gene expression heterogeneity is important to understand how these factors shape population structures and evolutionary adaptation, including drug resistance. We applied our scRNA-seq pipeline to generate a blueprint of single-cell transcriptional profiles in response to environmental changes, including stress agents, high cell density, or nutrient depletion, representing a total of 1824 cells (**Supplementary Table S1**). To compare and contrast single-cell responses to different environments, we focused on 110 genes that are upregulated as part of the CESR^29^. Transcriptional signatures resulting from different treatments could be clearly distinguished from our datasets using principal component analysis (PCA) of bayNorm normalised counts (Fig. 4a). Interestingly, cells growing rapidly in constant conditions occupied a distinct area of the transcriptional space, confirming our previous observation that exacerbated stress response is not a common feature of single cells during rapid proliferation (Fig. 2c). We then examined the specificity of the transcriptional programmes defined by scRNA-seq, restricting our analysis to genes specific for heat-shock and oxidative-stress responses^29^. Focusing on these specific gene sets singled out the cells that experienced the corresponding environmental stresses, and confirmed the specificity of scRNA-seq and the capacity of bayNorm normalisation to correct for experimental batch effects (Fig. 4b-c, **Supplementary Notes**).

**Figure 4:**
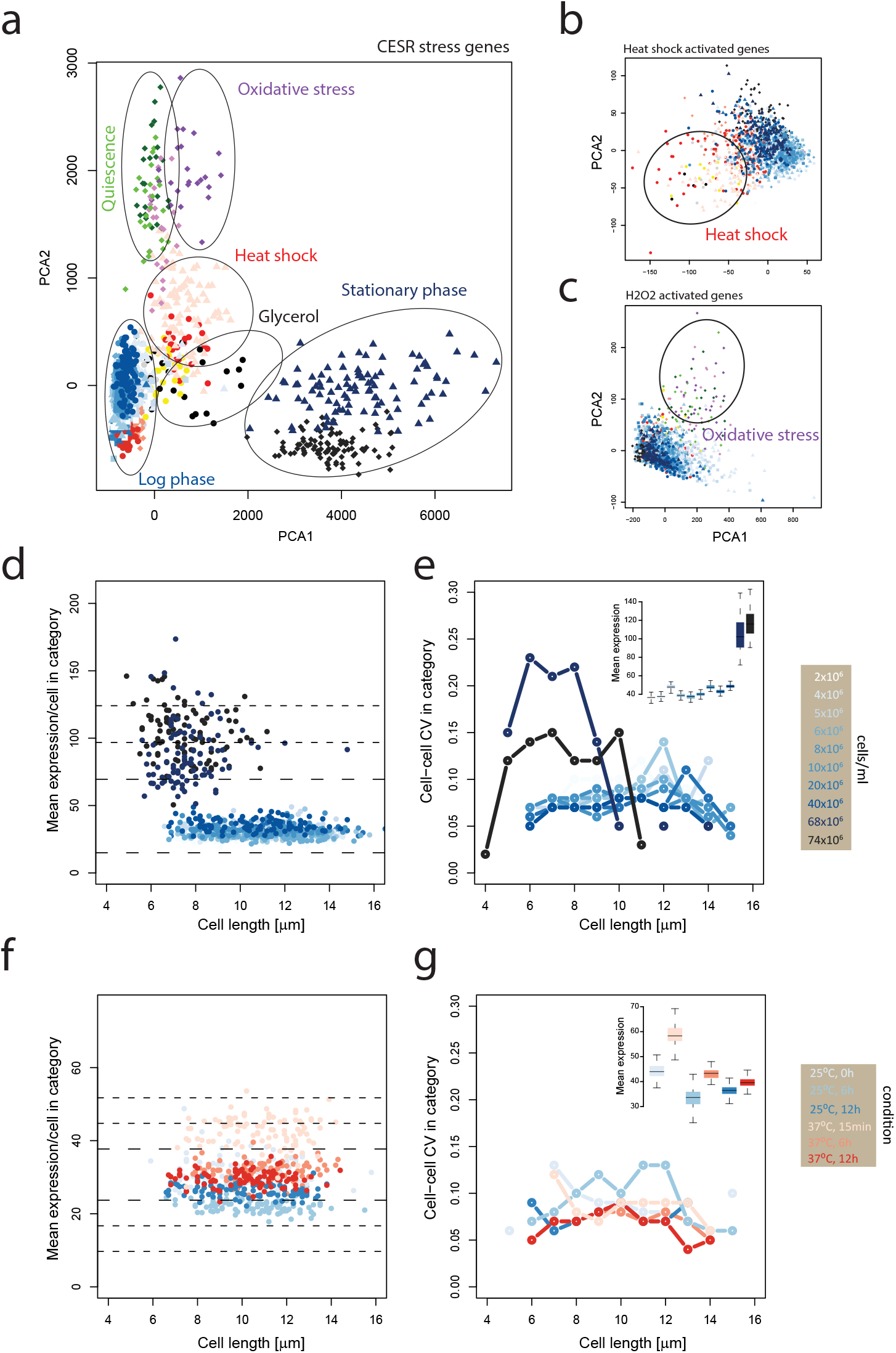
Gene-expression heterogeneity of fission yeast populations in response to environmental changes. **(a)** Single cells show distinct stress signatures of gene expression in response to different external conditions. Left: PCA analysis of normalised gene-expression scores for the core environmental stress response genes (CESR). A total of 1824 cells were analysed from cultures experiencing different external conditions (**Supplementary Table S1**). Each condition is colour coded, and larger groups are circled and annotated. **(b)** As in (a) showing genes specific for the heat-shock response^29^. **(c)** As in (a) showing genes specific for the oxidative-stress response^29^. **(d)** Heterogeneity in CESR gene expression during rapid proliferation and entry into stationary phase. Average CESR gene expression per cell plotted as a function of cell size. Cell density is colour coded as per the legend on the right. **(e)** Related to (d) showing between cell coefficients of variation in CESR gene expression (main panel) or average expression (insert). Note the strong increase in average CESR expression per cell accompanied by increased expression heterogeneity occurring at higher cell densities. **(f)** Heterogeneity in CESR gene expression during acute response and adaptation to heat shock. Average CESR gene expression per cell plotted as a function of cell size. Conditions are colour coded as per legend on right. **(g)** Related to (d) showing between cell coefficients of variation in CESR gene expression (main panel) or average expression (insert). Note the acute increase in average expression per cell of heat-shock genes and the lack of increase in expression heterogeneity during acute and adaptive responses.

We then asked whether the dynamics and heterogeneity of responses to environmental changes differed depending on the type of perturbation. We first analysed the gene-expression response of single cells to the gradual changes in external media conditions that accompany the increase in population density. To this end, we used scRNA-seq to analyse cells growing at densities ranging from 2×10^6^ to 74×10^6^ cells/ml, encompassing rapid proliferation and early stages of stationary phase (Fig. S4a, **Supplementary Table S1**). We observed a clear, progressive increase in concentration of CESR genes with cell density up to a concentration of ~40×10^6^ cells/ml (Fig. S4a-b). This increase in stress-related mRNAs was co-ordinated with a decrease in mRNAs participating in translation and the cell-growth programme (Fig. S4b)^29^. At the protein level, ribosomes have been shown to increase in concentration as a function of cellular growth rate in order to support the higher biosynthetic demand^46-48^. We therefore wondered whether the changes in mRNAs with cell density observed in our data happened in response to a change in cellular growth rates. Surprisingly, we found that population growth rates in our dataset remained constant up to densities of ~40×10^6^ cells/ml (Fig. S4a). This indicates that the decrease in mRNAs functioning in translation can take place in response to environmental changes independent of cell growth and without any immediate effect on growth. This observation is consistent with the existence of a free ribosome fraction that buffers growth and environment^48^. Importantly, although additional classes of mRNAs showed a similar coordination with cell density, this behaviour was not ubiquitous, suggesting that not all mRNAs respond to this form of environmental challenge (Fig. S4d, left panel). Notably, changes in concentrations of CESR mRNAs were neither accompanied by increased mRNA expression noise, nor by the appearance of cell outliers having entered a full stress-resistance state (Fig. 4e, S4d right). This result indicates that single cells perform a gradual and synchronised adaptation of gene expression to increased cell density before growth slows down during the transition to stationary phase (Fig. 4d).

Strikingly, within the following cell division, we detected a strong and heterogeneous increase in expression of CESR genes (Fig. 4d-e). This process coincided with the decrease in growth rate (Fig. S4a). Importantly, exhaustive functional analysis revealed that increase in transcriptional heterogeneity was restricted to specific pathways and not a global property of the transcriptome (Fig. S4d, **right**). Additional genes showing strong heterogeneous responses were also induced during meiotic differentiation and growth on glycerol (Fig. S4c **middle and right**)^49^. Taken together, these data support a model where single cells readjust the balance of the stress and growth transcriptional programmes synchronously as a function of cell density and ahead of changes in growth rate. This period is followed, within a single cell cycle, by a substantial, heterogeneous reshuffling of the cellular transcriptomes. These findings indicate that entry into stationary phase is a process that increases transcriptional heterogeneity and possibly promotes cell individuality and differentiation^50,51^.

In a second set of experiments, we examined the impact of a rapid and severe change in external conditions on gene expression at the cellular level. Cells were switched from 25°C to 37°C in a turbidostat and continued to grow at steady-state at either 37°C (adaptation) or 25°C (relaxation). We detected an increase in expression of the stress programme readily after the temperature switch, which adjusted back to pre-stress levels during both adaptation and relaxation (Fig 4f). Strikingly, we only detected a minor increase in transcriptional heterogeneity at the time of the heat shock that did not increase or propagate during adaptation or relaxation, in stark contrast with entry into stationary phase. Together this experiment suggests that an acute response to stress can be quite synchronous in a cell population and does not lead to phenotypic heterogeneity (Fig. 4g). Together, this analysis demonstrates that the level of transcriptional heterogeneity induced by changes in external conditions is variable and regulated, depending on the type of stimulus.

## Conclusion

We report a novel, integrated approach to analyse transcriptomes of single microbial cells in combination with phenotypic measurements. With this approach, we provide the first account of genome-wide heterogeneity of gene expression in fission yeast, both during rapid proliferation under constant conditions and in response to environmental changes. In constant conditions, periodic gene expression as a function of the cell cycle is the most robust and pervasive form of heterogeneity. Moreover, we report G2-specific expression signatures that overlap with signatures of the budding yeast YMC or that do not scale in concentration with cell size. This analysis was made possible by our ability to order and classify cells based on their length, independently of, but linked to, the scRNA-seq data. A setup for quantitative imaging of high-dimensional morphological features coupled to our approach would extend its potential even further, to more subtle cellular traits such as nuclear size, mitochondrial numbers, or actin structure. We also report the expression heterogeneity and its dynamics in response to environmental changes. We observed striking differences between entry into stationary phase, a heterogeneous process, and response to heat-shock, which appeared more coordinated. These differences highlight that expression heterogeneity is controlled in a condition-specific manner. An indepth analysis of a larger number of diverse environmental challenges will be required to understand the root of this variability in responses at the single cell level. In particular, it will be important to understand the impact of growth on expression heterogeneity and whether post-mitotic or quiescent cells that have exited the cell cycle show more homogeneous responses than cycling cells. Moreover, disentangling the respective contributions of transcriptional and post-transcriptional regulation to variable environmental responses will be key to reach a mechanistic understanding of this phenomenon. We show that a strong acute stress, such as a heat-shock, triggers a strong, homogeneous response. In this context, it will be important to understand how the heterogeneity of gene expression responses relates to the strength of the environmental challenge. Overall, in addition to increasing our understanding of how a simple eukaryote functions, the findings reported here highlight the power and potential of investigating gene regulation of unicellular organisms at both genome-wide and in single-cell levels.

## Methods

Detailed methods and associated references are available in the online version of the paper.

## Acknowledgements

We thank Tom Livermore for his help during the initial part of this project and Claire Stefanelli for her contribution to bayNorm development. We are grateful to Simona Parrinello and Amalia Martinez Segura for critical reading of the manuscript. This research was supported by the UK Medical Research Council, a Leverhulme Research Project Grant (RPG-2014-408), and a Wellcome Trust Senior Investigator Award to J.B. (grant 095598/Z/11/Z). It used the computing resources of the UK Medical Bioinformatics partnership (UK MED-BIO; aggregation, integration, visualisation and analysis of large, complex data), which is supported by the UK Medical Research Council (grant no. MR/L01632X/1) and the Imperial College High Performance Computing Service.(services/ict/self-service/research-support/hpc). All RNA-seq datasets are available in ArrayExpress accession number XXXXXXX.

---

**Figure S1:**
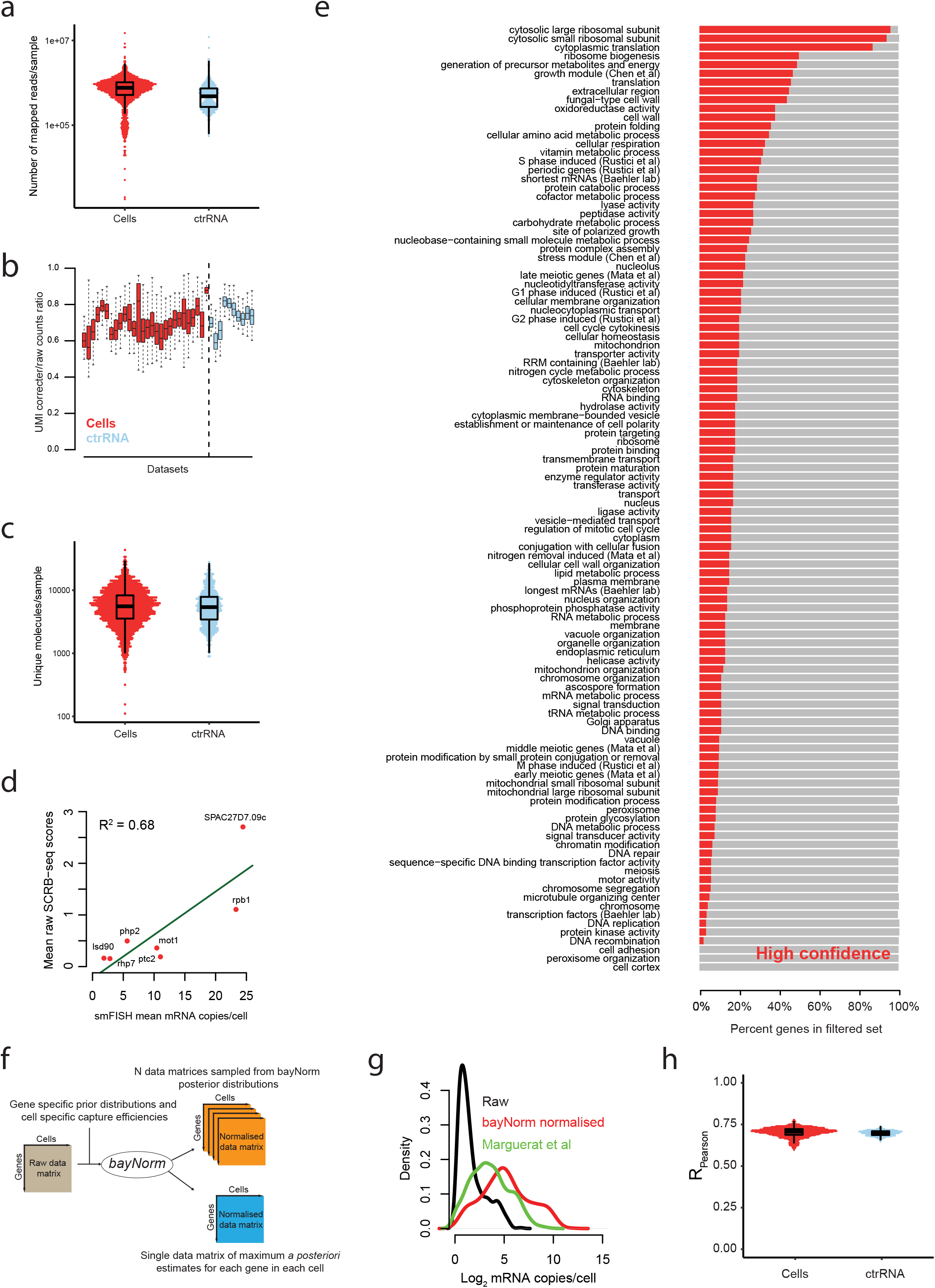
Supplement to *Imaging and transcriptome analysis of single fission yeast cells*. **(a)** Distribution of numbers of mapped reads per sample in cells (red) and ctrRNA (blue) datasets. **(b)** Fraction of molecules that passed filter after correction for UMI errors (Hamming distance < 2). **(c)** Distribution of unique mRNA molecule numbers per sample after UMI filtering in cells (red) and ctrRNA (blue) datasets. **(d)** Comparison of average bayNorm normalised values and average absolute mRNA counts defined by smFISH for seven genes (common names labelled on the plot). Regression line is shown in green and its R^2^ coefficient is shown in the top left corner. **(e)** Functional analysis of high-confidence gene dataset used in this study. The proportion of each gene category present in high-confidence dataset is indicated in red. Mean coverage across categories is 19.8%. **(f)** Cartoon of bayNorm normalisation procedure. Note the two different types of output that can be retrieved from the procedure. **(g)** Comparison of scRNA-seq raw and bayNorm counts averaged across cells with population estimates from Marguerat et al^21^. The shift between the green and red curves depends on the choice of 〈*β*〉. **(h)** Pearson correlation between bayNorm normalised molecule numbers/cell in individual cells (red) or ctrRNA (blue) datasets and population average mRNA copies/cells data from Marguerat et al^21^.

**Figure S2:**
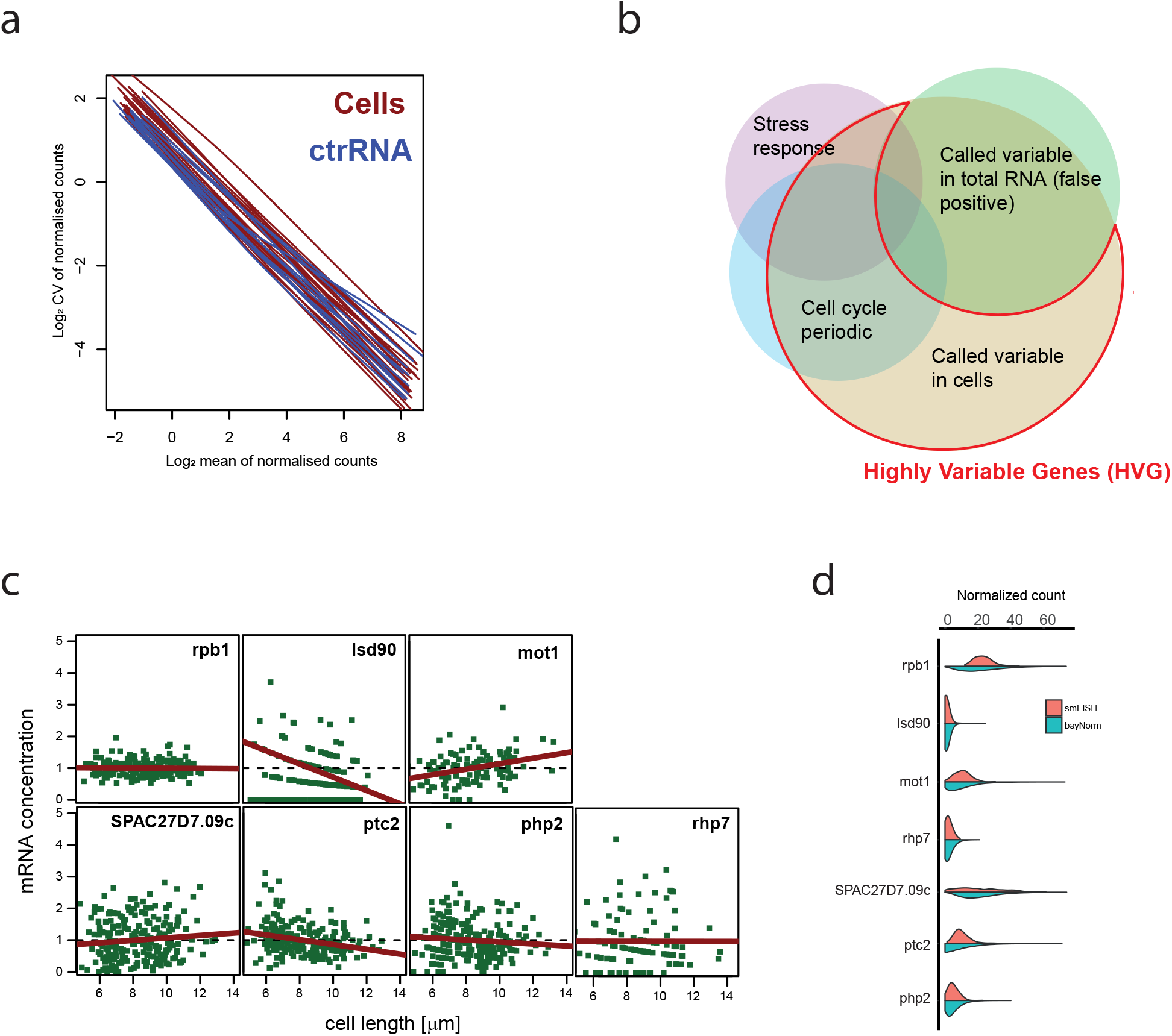
Supplement to *Cell-to-cell variability of the fission yeast transcriptome*. **(a)** Relation between mean expression and coefficient of variation (CV) for cell (red) and ctrRNA datasets (blue). Loess fits are plotted. Note the strong inverse relationship across all expression levels. **(b)** Identification of highly variable mRNAs. Variable mRNAs were called from 9 datasets of 96 cells, or 7 sets of total ctrRNA controls to define false positives. Genes called variable in at least 1 cell dataset, but never in ctrRNAs, constitute the HVG list used throughout this study (red circle, **Supplementary Table S2**). Note that mRNAs described as cell-cycle regulated or part of the core environmental stress response (CESR) show very low levels of false positives, thus validating the approach. **(c)** Singlemolecule fluorescence *in situ* hybridisation (smFISH) measurements of mRNA concentrations/cell plotted as a function of cell length. Regression lines are shown in red. **(d)** Comparison of smFISH molecule numbers and bayNorm normalised scores for 7 genes. Data were median centred to allow cross-comparison. Note the similar count distribution between both approaches.

**Figure S3:**
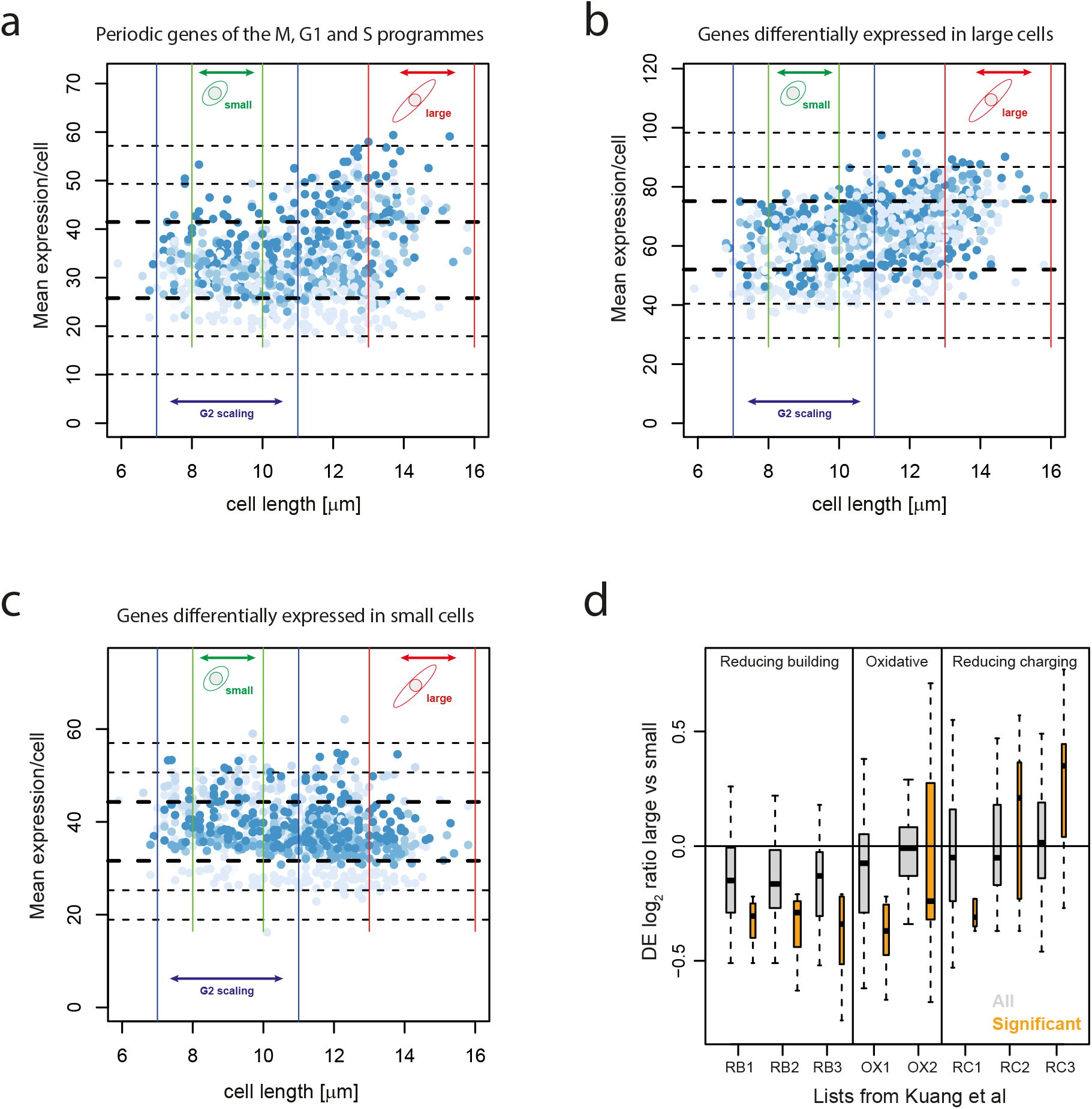
Supplement to *Cell-size dependence of fission yeast transcriptome*. **(a)** Mean expression per cell of genes associated with the M, G1 and S phases of the cell cycle^24^ as a function of cell length. Fission yeast cells during rapid proliferation are shown (n = 864). The size ranges of small (green bars) and large (red bars) cells used for differential analysis are shown. Blue bars show the size range used for defining non-scaling genes (NSG). **(b)** As in (a) for genes significantly up-regulated in large cells late in the cell cycle compared to small cells in G2. **(c)** As in (a) for genes significantly up-regulated in small cells in G2 compared to large cells late in the cell cycle. **(d)** Signatures of the Yeast Metabolic Cycle (YMC) in single fission yeast cells in asynchronous cultures. Differential gene expression log_2_ ratios between large and small cells were obtained with MAST and are shown for sets of genes participating in the YMC. Eight gene lists containing genes from the three phases of the YMC (top) were obtained from Kuang et al^52^. For each list, ratios of genes significantly regulated are shown in orange next to all genes in grey.

**Figure S4:**
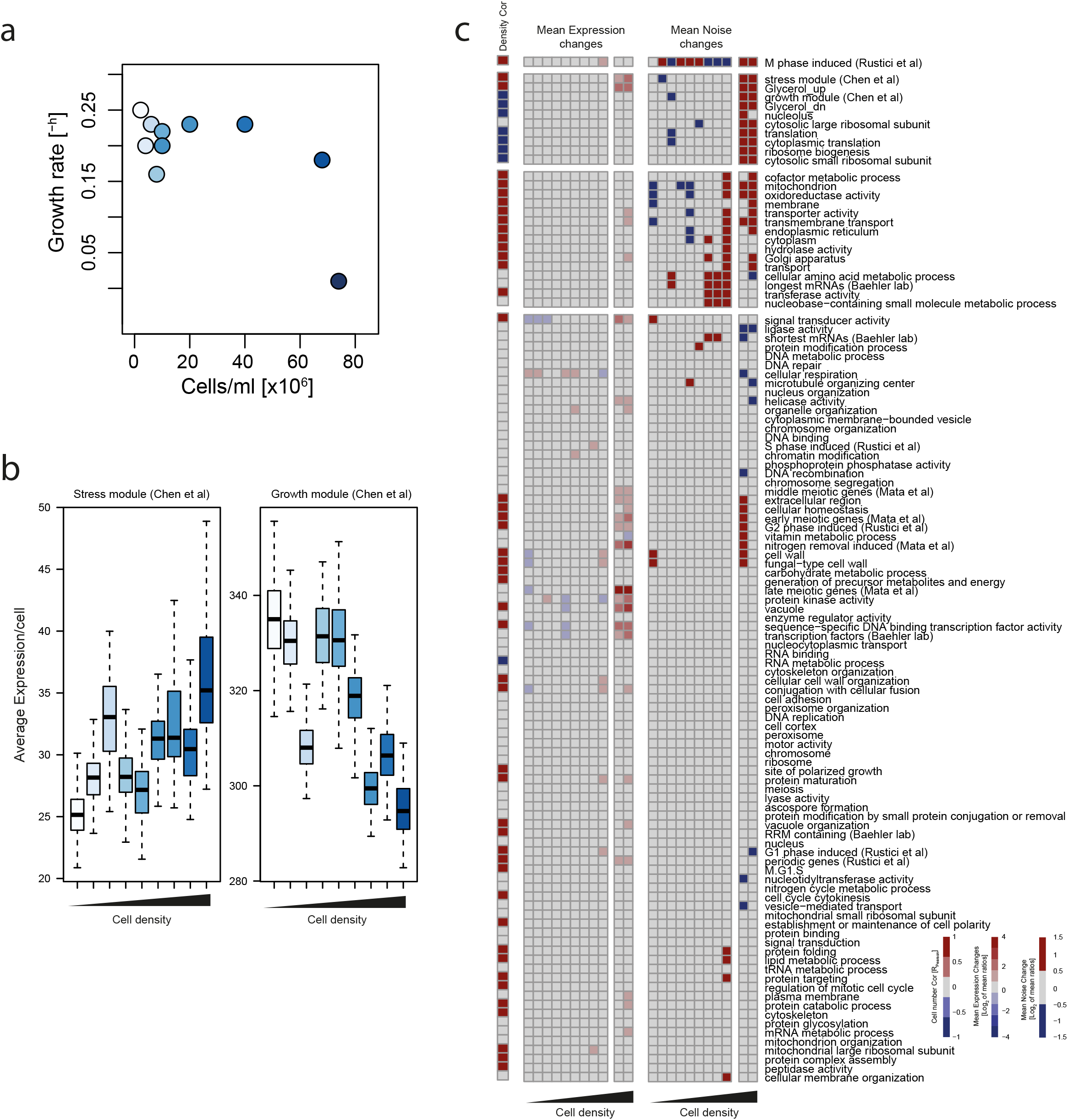
Supplement to *Gene-expression heterogeneity of fission yeast populations in response to environmental changes*. **(a)** Related to Figure 4d-e. Population growth rate plotted as a function of cell density. Conditions are colour coded as in Fig. 4d-e. The red dotted lines are a polynomial fit to the data to guide the eye. Note the constant growth rate up to a cell density of >44 × 10^6^ cells/ml. **(b)** Average expression per cell for genes from the stress response (left) and growth (right) programmes^29^. Boxes are colour-coded according to cell density as in Fig. 4d-e. **(c)** Functional analysis of gene expression changes during growth and entry into stationary phase. Left (Density Cor): Correlation of mean expression levels of gene categories with cell densities between 2-44 × 10^6^ cells/ml. Note that some categories increase in concentration co-ordinately with cell density, including the stress response programme, while other decrease, e.g. components of the ribosome, reminiscent of the P and R proteome fractions described in microorganisms^53^. Middle: changes in average expression of functional categories with cell density. For each category, the mean expression of genes from the category in each of the 11 cell-density datasets from Fig. 4d-e is compared to the other 10 datasets (10 ratio per density dataset). Log2 of the mean of the 10 ratios is shown for each density dataset. Right: as in middle but using noise measurements for each category and dataset. This analysis demonstrates that expression heterogeneity occurs during entry into stationary phase for specific categories only.

## Cell culture

972h- fission yeast cells were cultured with a seeding density of 0.5 × 10^6^ cells/ml in all experiments. Standard EMM2 media was used except indicated otherwise^1^. *Heat stress:* cells were grown in YE medium at 25°C up to a density of 2-4 × 10^6^ cells/ml. The cells were transferred to a water bath maintained at 37°C or 39°C (for datasets, 1712_1, 1712_2, 0408_2 (“Heat”)). For studying recovery from heat shock, cells were transferred post heat-shock to a turbidostat maintained at density of 4 × 10^6^ cells/ml and a temperature of either 25°C or 37°C (siphon-flow based derivative of the instrument described in ref ^2^). *Glycerol growth:* cells were grown in YE medium with 3% glycerol and 0.1% glucose. *Osmotic shock, Oxidative stress:* Cells were cultured in YE medium up to a density of 4 × 10^6^ cells/ml. To induce osmotic shock, an equal volume of 2M sorbitol prepared in YE medium to a final concentration of 1M sorbitol. To induce oxidative stress, cells were treated with 0.5mM H_2_O_2_ for 15 or 60 min. *Nitrogen starvation:* Cells were pre-cultured in EMM2 medium with NH_4_Cl as nitrogen source up to a density of 2 × 10^6^ cells/ml. Cells were harvested by centrifugation, washed twice with EMM medium without nitrogen and re-suspended in medium without nitrogen. Cells were harvested at 6 and 24 hours after starvation. All cultures were snap frozen at the time of harvesting.

## Cell isolation and imaging

Single cells were imaged with a 20x objective on a Singer MSM-400 tetrad-dissection microscope, picked into 3μl of QuickExtract™ for RNA extraction solution (Lucigen, Epicenter) in 200μl PCR tubes and immediately snap frozen at −80°C. The use of the QuickExtract buffer solution is critical for protecting RNA against degradation during cell lysis. For ctrRNA samples 3pg of total RNA isolated from matching culture as in were diluted in QuickExtract buffer solution and processed as single cells. Single cell images were analysed using ImageJ. All cells are described in **Supplementary Table S2**.

## SCRB-seq for yeast library preparation

Single cells were lysed at 98°C for 10 min in a PCR machine, and library preparation performed based on^3–5^, using primer sequences described in^3^. The protocol was modified as follows. Briefly, oligo dT containing cell indexes 1-96 and UMIs were added to each well at a final concentration of 1μM. Primers were annealed to the RNA template at 72°C for 3min, and components for reverse transcription added with final concentrations of 100U Superscript II reverse transcriptase (Invitrogen), 10U RNAse inhibitor (Invitrogen), 1x Superscript buffer, 5mM DTT, 1M Betaine (Sigma), 1.5mM MgCl2, 1mM of each dNTPs, 1/10^6^ dilution of ERCC spikes Set A (NEB), 1μM RNA-TSO primer. Reverse transcription was carried out at 42°C for 90min, after which the temperature was ramped between 50°C and 42°C for 10 cycles of 2min each. The reaction was heat inactivated at 70°C for 15min and the reaction cooled to 15°C. Each single cell sample was treated with 20U Exonuclease I (NEB) for 30min at 37°C followed by heat inactivation at 80°C for 20min. Sets of 96 samples were pooled and purified using a PCR purification kit (Qiagen) and eluted in 60μl elution buffer (EB) containing 10mM Tris-Cl, pH 7.5. Samples were treated with 40U of Exonuclease I for 30min at 37°C for a second time followed by heat inactivation at 80°C for 20min. PCR was performed on the pooled sample adding 1x KAPA HiFi buffer, 0.075mM of each dNTPs, 1μM PCR primer, and 1.25U KAPA HiFi enzyme. PCR cycling was done with denaturation at 98°C for 3min, followed by 25 cycles of denaturation, annealing and extension at 98°C, 60°C and 72°C for 20s, 15s and 1min respectively. A final extension of 72°C for 5min was done before cooling the samples at 15°C. Samples were purified using 0.6x Agencourt beads and eluted in 10-15μl of nuclease free 10mM Tris pH 7.5. Libraries were quantified on an Agilent Bioanalyser using an HS-DNA chip to confirm the presence of a clean peak at ~1000bp. Between 1-2ng of PCR library was used for tagmentation using the Illumina Nextera XT kit using a modified I5 primer as described in^34^. Between 8 and 12 PCR cycles were performed post tagmentation to amplify the 3’ fragments carrying the poly-A tail, the cell barcode and the UMI. The final libraries were purified using Agencourt beads twice at 1x bead concentration and final elution done in elution buffer. Libraries were quantified using on a Bioanalyser and sequenced.

## Single molecule fluorescence in situ hybridisation (smFISH)

Measures of cell size, mRNA number per cell and cellular mRNA concentrations where obtained for 7 genes by single molecule fluorescence in situ hybridisation (smFISH) as described in ref^6^. Genes are: SPBC16E9.16c (lsd90), SPAC27D7.09c, SPCC330.02 (rhp7), SPBC725.11c (php2), SPBC28F2.12 (rpb1), SPBC1826.01c (mot1), and SPCC1223.11 (ptc2). Processed data are provided in **Supplementary Table S6**. Probe sequences are provided in **Supplementary Table S7**.

## Sequencing and Read mapping

Pools of scRNA-seq libraries were sequenced on an Illumina HiSeq 2500 instrument at the MRC LMS genomics facility. Paired-end reads (150nt) were generated from two pools of 96 samples per sequencing lane. Data was processed using RTA version 1.18.64, with default filter and quality settings. The reads were de-multiplexed with CASAVA 1.8.4 (allowing 0 mismatches). Read1 was used to extract cell-specific indexes and Unique Molecular Identifiers (UMI). Corresponding Read2 were mapped to the fission yeast genome as described in ref ^7^. Mapped reads where assigned to fission yeast genes as described in ref ^7^ using Pombase annotation as of 27/05/2015 and including 5’ and 3’ UTR sequences. Read1 and Read2 were assigned to specific cell/RNA samples based on cell-specific indexes sequence de-multiplexed using in house Perl scripts. Within each specific cell/RNA sample, reads sharing identical UMI sequences and mapping to the same gene were collapsed.

## UMI Correction

Unique Molecular Identifiers (UMI) are short random DNA sequences, typically 6-8nt in length, which are appended to every single cDNA molecule during SCRB-seq for yeast sequencing library preparation. In SCRB-seq for yeast, UMIs are part of the first-strand reverse transcription primer^3^. UMIs are commonly used to remove PCR amplification biases but importantly have been recognized to be themselves prone to sequencing errors and biases^8–11^. These lead, for a given gene, to an enrichment in the fraction of UMIs with small sequence distances, also called Hamming distances that are higher than expected by chance^10^. This phenomenon is present in our data and leads to an overestimation of the library diversity. To correct for this bias, we developed an original network-based method which removes recursively, at each genomic loci, reads associated to UMIs that differ by a distance of 1 nucleotide (Hamming distance = 1) from the UMIs with the highest abundance. Our method is identical to the adjacency method introduced and implemented recently in UMI-tools^10^. Applying our UMI error correction method removes ~ 30% of the raw reads pool (Fig. S1b). For datasets description, statistics and raw counts see **Supplementary Table S1**, S2 and S3.

## Filtering out low confidence genes

Genes representing > 0.16% of the total molecules detected in the transcriptome of at least one cell across all datasets were included in the high confidence gene set used in this study. This empirical filter leads to a list of 1011 genes with varied functions and regulation patterns across conditions (Fig. 1b, Fig. S1e, **Supplementary Table S5**).

## Estimation of the mean capture efficiency using spike-ins and smFISH

We define the capture efficiency *β_i_* of a cell *i* to be the probability of observing (sequencing) any one of the cell’s original mRNA molecules. The mean capture efficiency 〈*β*〉 is the mean of the capture efficiencies across all cells. As, for any given gene, the observed UMI corrected counts per cell are lower than the original number of mRNA molecules present in a cell, capture efficiencies range between 0 and 1. Spike-in controls can be used to estimate *β_i_* and mean capture efficiency 〈*β*〉 ^12^. To do this, we divided the total number of spike-ins molecules observed within each cell by the corresponding theoretical number of input spike-ins molecules. The mean of these ratios across all the cells is 0.015. We believe this number is an underestimate of the true capture efficiency, given recent absolute estimates of average mRNA counts in fission yeast populations^7^. Consistent with our observations, it has been reported recently that spike-ins have a lower capture efficiency than mRNAs^11^. An alternative way to estimate mean capture efficiency relies on estimates of absolute mRNA molecules numbers per cell obtained by smFISH. Using 7 different genes, we fit a linear regression between the mean expression of UMI corrected sequencing counts and the mean of the corresponding smFISH counts. Using this approach, the mean capture efficiency is estimated to be the coefficient of variable, which is about 0.083 (Fig. S1d). In summary, the mean capture estimates obtained from spike-ins controls and smFISH measurements are very different. As a compromise, we chose to use the geometric mean of the two estimates, leading to a mean capture efficiency 〈*β*〉 of about 0.03. This estimate is one of the parameters for our Bayesian data normalization protocol described below (bayNorm)^13^. We note that, within this range, our biological conclusions are not overly sensitive to specific values of 〈*β*〉. The dependence of bayNorm normalization on the choice of 〈*β*〉 is systematically explored elsewhere^13^.

## Estimation of capture efficiencies of single cells

Single cells’ capture efficiencies (*β_i_*) are proportional to cell specific global scaling factors (*s_i_*) that are commonly used in normalization of single cell RNA-seq data (see for example^14^):

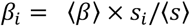

where, the constant of proportionality is related to the mean capture efficiency 〈*β*〉. The scaling factors (*s_i_*) can be estimated using spike-in controls^12^, or alternatively from the data directly. Simple estimates of *s_i_* are the total number of molecules detected per cell (total count, TC)^15^, or the mean of the number of molecules of a subset of genes detected in each cell. Popular bulk RNA-seq methods such as DESeq are designed to compute global scaling factors s_i_^16,17^. Unfortunately, these methods are not applicable to scRNA-seq datasets because of the high frequency of drop-outs present in the data (drop-outs: the proportion of genes with zero counts across cells)^14^. Alternative methods have been developed specifically for scRNA-seq (for example see^18,19^). We carefully assessed several existing methods for estimation of scaling factors, and settled for estimations of *s_i_* based on the mean of UMI corrected counts of a carefully chosen subset of genes in each given cell. The rationale behind this choice is the following: we argued that genes with i) high drop-out rates, ii) high variability in ctrRNA control experiments (showing technical variability) or iii) those in the tail of the mean expression distribution (which have disproportionally high contribution to the total count) are not suitable for scaling factor estimation. Specifically, we used a list of 768 genes for global scaling factors estimation that were selected as follows: i) Genes with dropout rates > 70% were excluded (zero UMI corrected counts in more than 70% of the cells across all datasets, 202 genes). ii) The top 20 higher expressed genes across datasets after TC normalization were removed. iii) Genes with significantly high technical variability in ctrRNA controls were removed. To do this we called highly variable genes in 11 ctrRNA datasets using the Bioconductor package scran and excluded 21 genes that were noisy in at least 5 of the 11 datasets^19,20^. Interestingly, the procedure described above produced the highest correlation between single-cell capture efficiencies and cell sizes (0.1781 for this procedure, 0.149 for the method proposed in^19^ and 0. 0568 for the spike-ins estimates).

## Normalisation using bayNorm

Single-cell RNA-seq data are commonly normalized by dividing raw counts by the global scaling factors *s_i_* estimated for each cell^14^. We have recently developed bayNorm an alternative Bayesian approach to scRNA-seq normalization that also provides simultaneous imputation for the drop-outs^13^. In this approach, given the raw count *x_ij_* observed in the *j^th^* cell for the *i^th^* gene and given the capture efficiency *β_i_* of the *j^th^* cell, we estimate the posterior distribution of the expected number of mRNAs 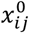 that was present originally in the cell. We found that a reasonable choice for the likelihood of observed counts *x_ij_* is a Binomial distribution with size 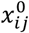 and probability *βi* ^13^. In addition, we assume that the prior for 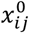 follows a Negative Binomial distribution (NB) with mean *μ_i_* and size factor *ϕ_i_*, with the following parameterization:

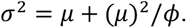

Using the Bayes rule, the posterior distribution of the *i^th^* gene in the *j^th^* cell can be expressed as:

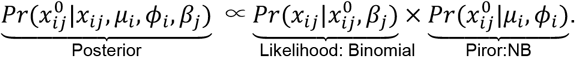

Outputs of the bayNorm normalisation procedure are either samples from the posterior distribution described above or its maximum *a posteriori* estimate (Fig. S1f). In addition to raw RNA-seq counts, bayNorm normalisation requires as inputs prior distributions of the parameters *μ_i_* and *φ_i_* for each gene. In bayNorm, these are estimated from the scRNA-seq data directly using an Empirical Bayes approach (see ref ^13^ for details). Prior estimation can be done using data from all cells across all datasets irrespective of experimental conditions. We refer to this procedure as “Global” normalisation. Alternatively, if cells can be split in different groups based on experimental conditions or phenotypic information for instance, prior parameters *μ_i_* and *φ_i_* can also be estimated within each group independently. We refer to this procedure as “Local” normalisation. On one hand, the use of global priors based on Empirical Bayes reduces strongly the technical batch effects that occur between different experiments. On the other hand, using local priors for different groups of cells enhances the resolution and sensitivity of differential expression analysis between these groups. Bayesian normalisation, as implemented in bayNorm, has several additional advantages over widely used normalisation approaches that rely on dividing molecules numbers by global scaling factors (see also^18^). First, bayNorm also provides imputation by replacing a large proportion of zero counts by non-zero values reducing strongly the fraction of drop-outs in the normalised data (from 42.27% to 3.56% in the cell datasets for the high confidence genes). Second, bayNorm effectively corrects for the experimental batch dependent variation in average capture efficiency and reduces batch-specific biases performing similarly to SCnorm but without the need of multiple expression dependent scaling factors^13,18^. Also, use of global priors as explained above can further reduce batch effects. Third, bayNorm normalisation preserves the uncertainty present in the data particularly for cells with low coverage, reducing false discovery rates in differential expression analysis. Finally, bayNorm produces mRNA distributions and noise estimates close to the state of the art smFISH measurements (Fig. S2g) and averaged transcriptomes structures close to high quality population estimates (Fig. S1g-h).

## Mean capture efficiencies, prior distributions, posterior distribution and point estimate datasets used in this study

Mean capture efficiency was set at 0.03 throughout the manuscript except for Fig. S2d where a capture efficiency of 0.08 calculated from smFISH data was used. Prior distributions where generated as follows: Fig. 1c, S1g, S1h, 3a, 3b, 3d, 3e, S3a-c, 4, S4 data were normalised using “global” priors obtained from all cells in the dataset in order to correct for batch effects. Fig S2d used priors calculated from rapidly growing cells. Fig. 2a, 2c, 2d, S2a, S2b data were normalised using “local” priors estimated within each individual datasets to exclude any residual contribution of batch differences to HVG calls. Figure 3c and S3d data were normalised using “local” priors estimated independently for sets of either large (13-16μm) or small (8-10μm) cells to maximize sensitivity of DE analysis.

## Detection of noisy or highly variable genes (Fig. 2)

Expression of a given gene can vary among cells of a population. Cell-to-cell variability in gene expression, also called noise, is defined as the coefficient of variation (*σ/μ*)^2^, where σ and μ are the standard deviation and the mean of expression scores across cells respectively. A number of modelling and experimental studies have shown that gene expression noise is inversely correlated with mean gene expression calculated across cells^21–24^. Genes with particularly high cell-to-cell variability are called noisy or highly variable genes (HVGs) and are defined as having significantly higher noise than most genes of similar means (Fig. 2a). Identifying HVGs from scRNA-seq experiments is challenging due to the strong technical noise present in the data and several teams have addressed this problem^20,23–25^. The general consensus is to decompose the total noise observed into its technical and biological components. To do this, the dependence of technical noise to the mean is measured and used to infer potential additional biological noise present for each gene (Fig. 2a, Fig. S2a). Here, we have developed an original method for HVG detection based on bayNorm normalised data and sets of computed synthetic control RNA-seq data (synRNA) to estimate noise floors. synRNA are generated as follows: Given a scRNA-seq dataset with a raw count matrix *x_ij_* and a vector of estimated cell-specific capture efficiencies *β_j_*, we produced a set of synRNA data 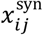 with similar mean expressions and capture efficiencies as the real experimental data but with no biological variability above what is expected from the Poisson distribution. Poisson noise is the minimal amount of noise expected if no additional biological variability is present. To do this, we first generate a gene expression dataset 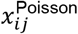 sampled from a Poisson distribution with mean expression obtained from raw count matrix *x_ij_* and capture efficiencies *β_j_*:

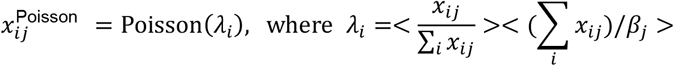

both means above are calculated as across cells (index *j*). In a final step, we used binomial downsampling to generate a 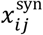 dataset from 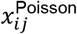 simulating the effect of partial RNA capture during the scRNA-seq procedure:

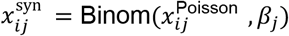

Finally, 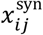 data are normalised with bayNorm using prior parameters estimated from the original raw data (i.e. identical to those used for normalisation of the cells data). To identify highly variable genes, a local regression between noise and mean expression of all genes from the normalized synRNA dataset is calculated and compared to the noise levels observed in the corresponding normalised experimental datasets (log-log, Fig. 2a). To call genes with noise levels significantly above the synRNA fitted line, we use an approach similar to the one proposed in ref ^20^ and based on an adaptation of the gene expression variation model^25^. Briefly, vertical differences (only considering positive residuals) between noise levels in the experimental dataset and the fitted line were calculated (illustrated as ΔCV on Fig. 2a). The differences were normalised by dividing by the residuals from the regression, which follow a normal distribution. Most genes were assumed not to deviate from the centre of the distribution significantly. The centre was found by the kernel density of the normalized differences. Next a normal distribution was fitted using differences which were below that centre. P-values were then extracted based on the normal distribution and adjusted using Benjamini and Hochberg^26^. Noisy genes were called independently for each batch of 96 cells or ctrRNAs generated in this study after normalisation with bayNorm using priors estimated within each dataset (“local” priors). For each gene in each dataset noise and mean values across cells where calculated using pooled expression scores of 100 samples of the bayNorm posterior distribution per cell. Using this design gene variability was assessed in 100 bootstrapped versions of each dataset. Genes were called noisy if they had an FDR < 0.1 in 85 or more bootstrap samples. Genes called in at least one rapid growth cell datasets (2502_1, 2502_3, 2502_5, 2502_7, 2502_9, 1904_1, 0109_3, 1711_1, 1711_2), and none of the ctrRNA datasets (2502_2, 2502_4, 2502_6, 2502_8, 2502_1, 1904_2, 0408_5), were called HVGs and discussed further in this study (**Supplementary Table S5**). Transcriptome fractions of functional categories (Fig. 3b)

To assign cells to particular functional categories, we calculated z-scores for the sums of the counts of each category within each cell. Cells were assigned to a given category if the category |z-score| was > 1.2 in more than 70 out of 100 bayNorm posteriors samples. Categories with assigned cells significantly larger or smaller than the whole population are shown on Fig. 3b (P_Wilcox_ < 0.05).

## Differential Gene Expression analysis (Fig. 3c, S3b-d)

Several Differential Expression (DE) analysis methods, tailored for scRNA-seq analysis have been published. A recent comparative analysis^27^ and our own experience identifies MAST^27^ as a reliable method. Therefore, in this study, we used the MAST package with method = “glm”, the “ebayes” option enabled, and considering adjusted P-values from the continuous part of the hurdle model utilized in MAST (multiple testing adjustment method: Benjamini and Hochberg)^26,28^. DE detection was run independently on 100 bayNorm posterior distribution samples for Fig. 3. Genes called DE in > 90% of the posterior distributions were considered differentially expressed. Log_2_ ratios are the mean of the log2 ratios from each posterior distribution. An additional cut off on log2 ratios (> 0.2 or < −0.2) was used on Fig. 3. Figure 3 DE analysis used two sets of cells either large (13-16μm) or small (8-10μm).

## Random forest model

To identify genes that have non-linear correlation of expression levels with cell size, we built a random forest model of cell size given gene expression levels. We chose a subset of normal cells and applied a filtering criterion such that cells with total counts less than 100000 or greater than 300000 were removed. In addition, we filtered out cells with length < 6μm or > 25μm. We then applied a Random Forest model described in^29^ on the filtered dataset. Genes were ranked according to the importance statistic “%IncMSE” returned by the model (**Supplementary Table S5**).

